# The Area-law of Molecular Entropy: Moving beyond Harmonic Approximation

**DOI:** 10.1101/2024.03.16.585357

**Authors:** Amitava Roy, Vishwesh Venkatraman, Tibra Ali

## Abstract

Inspired by black hole thermodynamics, the area law that entropy is proportional to horizon area has been proposed in quantum entanglement entropy and has largely maintained its validity. This article shows that the area law is also valid for the thermodynamic entropy of molecules. We showed that the gas-phase entropy of molecules obeys the area law with our proposed correction for the different curvatures of the molecular surface. The coefficient for the ultraviolet cutoff for the molecular entropy, calculated from our curated experimental data, is tantalizingly close to the value 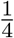 proposed by Hawking [Hawking, 1976]. The ability to estimate gas-phase entropy by the area law also allows us to calculate molecular entropy faster and more accurately than currently popular methods of estimating molecular entropy with harmonic oscillator approximation. The speed and accuracy of our method will open up new possibilities for the explicit inclusion of entropy in computational biology methods, such as virtual screening applications.

## 1 Introduction

Free energy governs all chemical processes. Change in free energy in a chemical process can be written as Δ*G* = Δ*H* −*T* Δ*S*_th_, where *G* is the Gibbs free energy, *H* is the enthalpy, *S*_th_ is the thermodynamic entropy governed by the second law of thermodynamics.

The second law of thermodynamics, as defined by Clasius [Clausius, 1879], can be stated as follows: without outside intervention, heat flows from hot to cold as a non-equilibrium system reaches equilibrium. Ludwig Boltzmann connected the macroscopic definition of entropy from Clausius’s definition with the microscopic property of a system by the celebrated equation

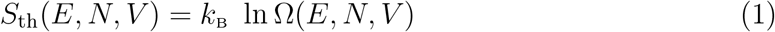

where Ω is the total number of distinct microstates accessible to the system with the given macroscopic constraints. From the postulation that a thermodynamic equilibrium for an isolated system is the state of maximum entropy, Gibbs arrived at a similar form for the entropy of a system. However, the two methods differ in calculating the number of microstates and can provide different entropy values for the same system [Jaynes, 1965]. Following Gibbss’ formulation, thermodynamical entropy (also often referred to as Boltzmann-Gibbs entropy) can also be expressed by

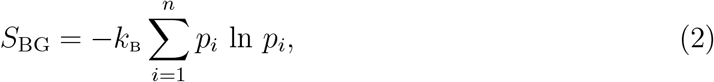

where *k*_B_ is the Boltzmann constant, *p*_i_ is the probability for the microstate *i*, and *n* is the number of microstates. Equation 2 is similar in form to Shannon’s formulation of entropy of information [Shannon, 1948],

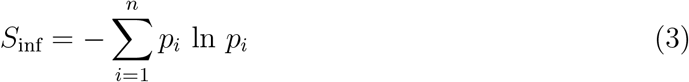

where *p*_i_ is the probability of the event *i*, and *n* is the number of events. However, one must be careful to interpret any term defined by equation 3 as thermodynamic entropy. Such an association has to explain the experimental observable of the heat flow from a hot to a cold body determined by the second law of thermodynamics.

The most noted work in connecting information-theory entropy with thermodynamic entropy was by ET Jaynes in his seminal paper “Information theory and statistical mechanics” [Jaynes, 1957]. Jaynes has argued that for a given macroscopic constraint, the randomness of the microstates can be estimated by information theory in the least biased way. Furthermore, the maximum-entropy principle is sufficient to develop the rules of statistical mechanics, from which the equilibrium thermodynamics properties can be calculated [Jaynes, 1957]. Jaynes’s maximum-entropy principle states that given precisely stated prior data, the probability distribution with the largest entropy best represents the current state of knowledge about a system [Rosenkrantz, 1989].

### Computational techniques to calculate the entropy of a single molecule

Temperature is not well defined for a single molecule; consequently, the entropy of a single molecule cannot be understood in terms of heat flowing from a hot body to a cold one. Instead, we will use a definition of entropy similar to the one used by Jaynes - entropy represents uncertainty in our current state of knowledge or missing information (MI) (a term coined by Ben-Naim [Ben-Naim, 2008]) of a system under given macroscopic constraints. This article will refer to the entropy of a single molecule as *S*_mol_.

The most common approach for calculating the entropy of a single molecule is to approximate the dynamics of the molecule using Born–Oppenheimer approximation - the motion of an isolated molecule can be approximated by the motion of the nuclei in the potential created by the surrounding electrons. The approximation allows us to write molecular entropy as a sum of positional, orientational, and vibrational entropy, assuming that a molecule’s positional and orientational entropies are not coupled with the vibrational entropy of the molecule. However, orientational entropy is not necessarily decoupled from vibrational entropy for a flexible molecule ([Littlejohn and Reinsch, 1997]). Note that positional and orientational entropies are also referred to as translational and rotational entropies.

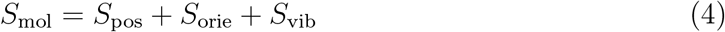

Vibrational entropy can be divided into harmonic, anharmonic, and *configurational* entropies (*S*_conf_), especially for flexible and complicated molecules.

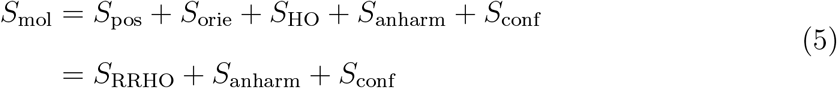

*S*_HO_ denotes the entropy of a harmonic oscillator, *S*_anharm_ is the anharmonic correction, and *S*_conf_ is the configurational entropy arises when a molecule can exist in multiple stable conformations at a given macroscopic constraint. Different conformations’ vibrational entropy must be appropriately averaged for a molecule with multiple stable conformations to calculate configurational entropy [Karplus et al., 1987]. Note that, in the literature, the terms configurational and conformational entropies are sometimes used interchangeably. Explicit calculations of *S*_anharm_ are often neglected or absorbed in the calculation of *S*_conf_. Calculating *S*_conf_ is intrinsically computationally expensive, as it requires, in principle, sampling the entire phase space. Consequently, calculating accurate entropy for a molecule can take thousands of CPU hours using molecular dynamics with molecular mechanics approximation (approximating the quantum mechanical interactions with a classical mechanical model [Van Gunsteren et al., 1994]). Please see [Chan et al., 2021b] for a recent effort to calculate *S*_conf_ for small molecules using molecular dynamics simulations.

One way to reduce computational resources is to approximate a molecule as a collection of simple harmonic oscillators (SHM), use a normal mode analysis technique (NMA), and calculate vibrational entropy. This approximation allows for avoiding identifying and counting microstates. The density matrix of a collection of SHMs is generally simpler to form in the energy-eigenstates, eigenstates of the Hamiltonian, basis from which entropy can be calculated, avoiding identifying and counting microstates. However, this approximation of expressing the vibration of molecules as a collection of harmonic oscillators is not appropriate, as anharmonic vibrations play an equally important role in the dynamics of small molecules, as do harmonic vibrations [Christiansen, 2012]. Still, NMA calculations help examine a molecule’s dynamics [Ma, 2005]. Vibrational entropy is replaced by configurational entropy when a molecule can exist in multiple stable conformations at a given macroscopic constraint.

Calculating molecular entropy requires the calculation of orientational and positional entropy along with vibrational or configurational entropy. We can easily calculate molecules’ orientational and positional entropy from their geometry if we approximate them as rigid bodies. Please see [Murray and Verdonk, 2002] for a review of the topic. The entropy approximated by the rigid body and SHM approximation is referred to as *S*_RRHO_ (eqn. 5) - rigid rotor and harmonic oscillator approximation. To calculate the entropy of a molecule using SHM approximation, charge distribution and the molecule’s geometry must be known. If the charge distribution is calculated using molecular mechanics approximation, the entropy can be computed using CPU-minutes to CPU-hours of computational resources. If the charge distribution is calculated using quantum mechanical methods, it may take CPU-days to CPU-weeks of computational resources.

In this article, we propose an alternate way of calculating molecular entropy calculated from the surface properties of a molecule. The foundation of our approach is motivated by the area law of entropy.

### Area law - an alternate way of calculating entropy

Bekenstein, in his celebrated paper, proposed that the thermodynamical entropy of a black hole is proportional to its surface area [Bekenstein, 1973]. The proposal is counter-intuitive as one would expect that the volume of the matter contains information; consequently, entropy should scale with volume but not surface area. In the article, Bekenstein explains how the counter-intuitive proposal is grounded in a deeper consideration of physics. At the same time, Hawking also derived a simple formula for the black hole’s entropy, equivalent to Bekenstein’s equation for entropy [Hawking, 1976]

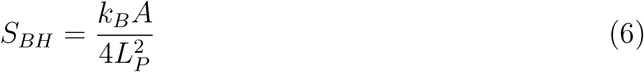

where *A* is the area of the horizon of the black hole, *k*_B_ is the Boltzmann constant, and *L*_P_ is the Planck length, the minimal value of the length used in the derivation (the ultraviolet cutoff *L*_UV_). Hawking also showed that the area of the horizon of a classical black hole never decreases. As a consequence, *S*_BH_ of a classical black hole never decreases. The postulate that the total horizon area of classical black holes cannot decrease was recently proven experimentally by Laser Interferometer Gravitational-wave Observatory (LIGO) [Isi et al., 2021].

The intriguing connection of black hole entropy with its surface area gave rise to several hypotheses on whether black hole entropy counts the microstate entropy and whether the relationship between entropy and area is a fundamental aspect of nature. For a review of the field, please check the reference [Eisert et al., 2008]. Early seminal works identified quantum entanglement entropy as also proportional to area [Bombelli et al., 1986, Srednicki, 1993]. (See Supporting Information for a brief introduction to quantum entropy and quantum entanglement entropy). Srednicki, in his work, showed that if a system of coupled harmonic oscillators is divided into two regions, oscillators inside an imaginary sphere *I* and oscillators outside the sphere *O*, and the density matrix is traced over the *O* oscillators, the entanglement entropy of the reduced density matrix is proportional to the area of the sphere that encloses *O* [Srednicki, 1993].

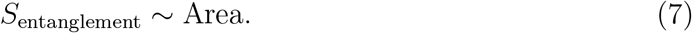

For the derivation, Srednicki chose a local Hamiltonian, i.e. the entanglement between the oscillators, which lie far from each other, contributes very little to the entanglement entropy. Intuitively, for a quantum system with many microscopic degrees of freedom, the significant contribution to the bipartite entanglement entropy comes from the entanglement of the states of the degrees of freedom that lie near the boundary.

It is impossible to probe the area law of entanglement entropy with current experimental advancement. The reference [Herdman et al., 2017] provides a detailed simulation study of superfluid ^4^He where the area law of entropy has been verified directly.

### Thermal entropy from quantum entanglement

The discussion of the area law of quantum entanglement has so far been for systems at zero temperature or in the ground state. When we measure the energy of a subsystem of a system in the ground state, there will be particular uncertainty of the energy of the subsystem from the quantum entanglement. The observed energy value will be a spread of energy with a peak at the expected ground-state subsystem energy value. This is readily seen from the reduced density matrix being in a probabilistic mixture of different energy eigenstates. Suppose we change this subsystem’s energy very quickly, something known as quenching, and let the entire system become thermally stable. Given a sufficient number of degrees of freedom, the system will become approximately a thermodynamic ensemble. In such a scenario, it has been theoretically predicted [Deutsch et al., 2013, Santos et al., 2012, Garrison and Grover, 2018] and experimentally shown [Kaufman et al., 2016] that the expectation value of any observable arising due to the uncertainty of quantum entanglement entropy at zero degree temperature and due to the thermodynamic entropy of the entire system at a finite temperature is identical. It is hypothesized that the signature of thermodynamic uncertainty of the full system is embedded in the quantum entanglement entropy of the subsystem.

Motivated by the fact that our uncertainty in knowledge about a system at a finite temperature due to the thermodynamic entropy is identical to the uncertainty arising from quantum entanglement entropy, and quantum entanglement entropy is proportional to the area of the system, we propose a method to estimate thermodynamical entropy of a molecule from its surface property as described below.

### Estimating molecular entropy using the area law

Embracing the area law of entropy from quantum entanglement, we postulate that the thermodynamic entropy, i.e., given macroscopic constraints like pressure, volume, temperature, and number of atoms, the uncertainty in the microstates of a molecule is proportional to its surface area. The area law developed for the different systems, from black holes to simple harmonic oscillators to ^4^He ions, is for primarily spherical surfaces and occasionally for other regular geometric surfaces. Most molecular surfaces are not of any regular geometric shapes and contain various degrees of curvature, which can be measured by a molecular surface’s shape index (𝒮) (see Method) values. As the effect of the curvature in entanglement entropy has yet to be studied in detail, we make some assumptions for simplicity.

1. Surface deformations, curvature with a 𝒮 value other than 1, change the uncertainty of our knowledge about the system’s microstates. Note that 𝒮 value 1 indicates a perfectly spherical surface.
2. Positive (0 ≤ 𝒮 *<* 1) and negative (-1 ≤ 𝒮 *<* 0) deformations change the uncertainties our knowledge about the system independently.

We can express the degree of uncertainty of our knowledge about the molecular system due to the deformations with Shannon’s formulation of entropy of information (Equation 3). Furthermore, following Jayne’s work [Jaynes, 1957], we can express thermodynamic entropy in terms of the uncertainty of our knowledge about the system. Consequently, we can write the thermodynamic entropy of a molecule as

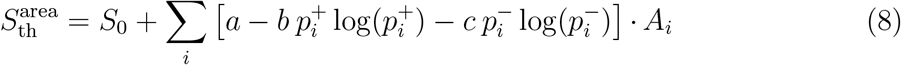

where, 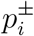 is the probability that the *i*th surface patch, with area *A*_i_, will have a shape specific positive or negative value of 𝒮. The constants *S*_0_, *a, b*, and *c* can be temperature dependent, where *S*_0_ has the dimension of entropy, and the rest has the dimension of entropy per unit area. The deformations, i.e., deviation of shape index value from 1, arise when the surfaces of more than one atom overlap. Such deformations indicate atomic bonds. As the number of bonds in a molecule increases, the uncertainty of our knowledge about the system’s microstates decreases. Consequently, we expect the overall signs of the terms related to the deformations to be negative. The individual *p log p* terms are all negative, and the constants *b* and *c* should be negative to make the deformation terms in our equation negative. Ideally, the constants *S*_0_, *a, b*, and *c* should be derived from an *ab initio* model of the area law of the molecular entropy. In this article, we collected experimental values of gas-phase entropy of 1942 small molecules and fit the data to derive the constants. As the collection of molecules increases, the constants from fitting the experimental data should approach the universal constants.

The deformations, i.e., deviation of shape index value from 1, arise when the surfaces of more than one atom overlap. Such deformations indicate atomic bonds. As the number of bonds in a molecule increases, the uncertainty of our knowledge, or the MI, about the system’s microstates decreases. Different deformations values, or 𝒮_i_, may represent specific arrangements of chemical bonds, and the probability *p*_i_ associated with 𝒮_i_ may reduce the MI of the system in a particular manner which is universal. If that is true, we ideally should calculate the *p*_i_ associated with 𝒮_i_ from the surface features of all possible molecules, which is practically impossible, or from the *ab initio* derivation of the *p*_i_s, which not in the scope of the current article. In this article, for practical purposes, we calculated the values of 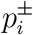 independently for each molecule under study separately.

## 2 Results and Discussion

From the literature, we curated 1942 experimental gas-phase entropy values of organic molecules, i.e., involving elements C, H, N, O, S, P, Cl, Br, and I. Gas-phase entropy is expressed as the molar entropy or entropy content of one mole of pure gas at standard pressure and temperature. Gas-phase entropy can be measured by measuring a gas’s heat capacity as a function of temperature. Please see [Kabo et al., 2019] for a review of the experimental methods for measuring the thermodynamic properties of organic substances.

For the curated 1942 molecules, most of the *S*_Expt_ values were between 250–500 *J/mol* · *K*, and entropies typically increased with the number of atoms in the molecule (Fig. S1 in SI). We calculated the structures of the 1942 molecules from their SMILES strings, and from the structures, we calculated the molecular surfaces and shape index values. Please see the Methods section for details of each of the steps. In subsequent sections, we show that the thermodynamical entropy calculated from a molecule’s area and shape index values has a collinear relationship with the gas-phase entropy. From the collinearity, we can estimate the gas phase of a molecule in a matter of seconds compared to tens to hundreds of CPU hours for other methods.

### Gas-phase entropy varies linearly with shape entropy

In our data set, the thermodynamical entropy 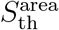 calculated from the surface features of a molecule showed a linear relationship with experimental gas-phase entropy, *S*_Expt_ (Fig. 1), with *R*^2^ values of ≈ 0.98 (Table 1). Note that, in this article, the area is measured in Å^2^. The root mean square error (RMSE) with the experimental values is 21.32 *J/mol* · *K*, and the mean average percentage error (MAPE) is 3.94 %. The *b* and *c* in Equation 8, the coefficients for the deformations, are as expected negative and -6.910 and -21.456, respectively. The deformations indicate atomic bonds and consequently reduce the uncertainty of our knowledge about a molecular system’s microstates. The linear relationship holds for the full range of the experimental gas-phase entropy values: 190 – 1040 *J/mol* · *K*. Even if the effect of the deformation is not taken into account, and the entropy is modeled as proportional to the surface area, the RMSE with the experimental values is 27.97 *J/mol* · *K*, and MAPE is 5.42 % (Table 1). We calculated the 95% confidence intervals using bootstrapping [Chernick, 2007] for each parameter.

**Table 1:** Root mean square error (RMSE), coefficient of determination (*R*^2^), and mean average percentage error (MAPE values) between predicted and experimental gas-phase entropy. In *S*_area_, we assumed the entropy is proportional to the surface area and ignored any contribution from the deformations in the surface. In this article, the area is measured in Å^2^. In 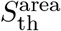, we added the contribution of the deformations. In G4 and NMA, entropy values were calculated using rigid rotor harmonic oscillator (RRHO) approximation, where the vibrational entropies were calculated using density functional theory and normal mode analysis (NMA) using molecular mechanics forcefield, respectively. We calculated *S*_area_ and 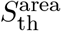 for all 1942 molecules, and the only parameters used were the atomic van-der waals radii of the atoms from OpenBabel [O’Boyle et al., 2011], which calculates a single VDW radius for each element regardless of its environment using the method described in [Mantina et al., 2009]. For G4, we curated and calculated entropies for 1529 molecules. Calculations for the rest of the molecules did not converge in our stipulated time frame. For NMA, we calculated entropies for 1665 molecules. We could not confidently generate the forcefield parameters for the rest of the molecules. We used bootstrap methods (100 bootstraps) to calculate lower and upper limits of 95% confidence interval. Upper and lower limits are shown as raised and lowered numbers, respectively.

| Method | $S_{\text{gasphase}} =$<br>$(J/mol \cdot K)$ | RMSE<br>$(J/mol \cdot K)$ | $R^2$ | MAPE<br>% |
| --- | --- | --- | --- | --- |
| Area | $S_{\text{area}} = 118.84_{118.62}^{119.23} + \sum_i 1.948_{1.945}^{1.959} A_i$ | $27.977_{27.841}^{28.112}$ | $0.957_{0.955}^{0.958}$ | 5.42 |
| Area +<br>deformation | $S_{\text{th}}^{\text{area}} = 104.32_{104.06}^{104.47} +$<br>$\sum_i [3.078_{3.068}^{3.083} + 6.910_{6.819}^{6.980} p_i^+ \log(p_i^+) +$<br>$+ 21.456_{21.352}^{21.726} p_i^- \log(p_i^-)] \cdot A_i$ | $21.326_{21.281}^{21.542}$ | $0.975_{0.974}^{0.976}$ | 3.94 |
| G4 | $S_{\text{RRHO}} = S_{\text{pos}} + S_{\text{orie}} + S_{\text{HO}}$ | $24.762_{23.180}^{26.480}$ | $0.945_{0.937}^{0.953}$ | 4.09 |
| NMA | | $45.441_{42.86}^{48.51}$ | $0.971_{0.976}^{0.967}$ | 5.41 |

**Figure 1:**
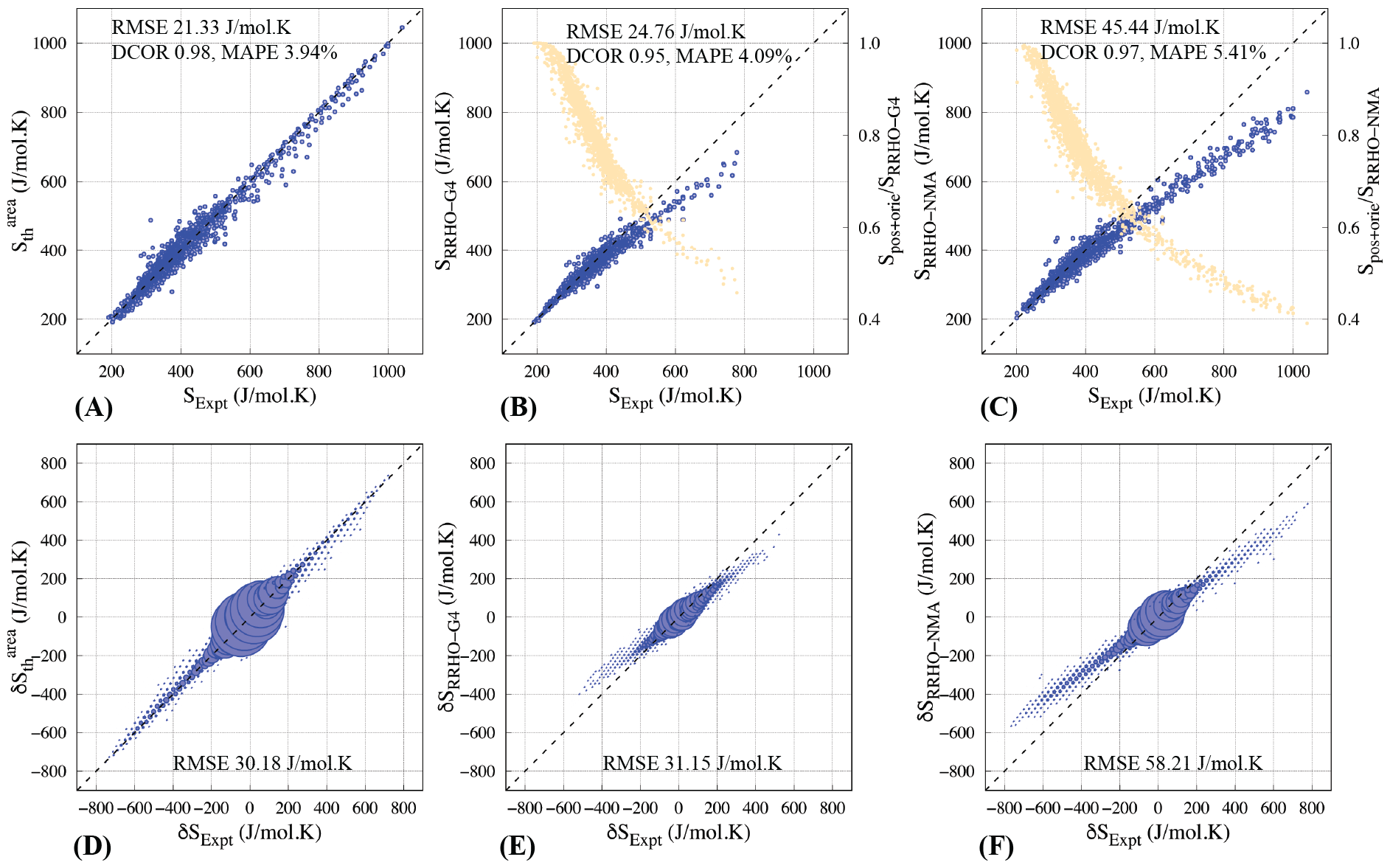
(A) 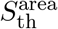, the thermodynamical entropies calculated from the area law (eqn. 8) are plotted against experimental gas-phase entropies for 1942 molecules. The root mean square error (RMSE) between the calculated and experimental entropy is 21.34 *J/mol* · *K*. The correlation (DCOR) between the values, calculated using distance correlation (see Methods), is 0.97, and the mean average percentage error (MAPE) is 3.94%. The dotted line represents the line where the values of the X and Y axes are equal. (B) Thermodynamic entropies were calculated using G4 quantum chemical calculations with the SHM approximation and plotted against experimental gas-phase entropies for 1529 molecules (blue dots). For the remaining 413 molecules, mainly the larger molecules, the G4 calculations did not converge. The orange dots represent the positional and orientational entropy as a fraction of the calculated total entropy. The error in the calculated entropy increases as the positional and orientational entropy falls below 60% of the calculated total entropy. (C) Thermodynamic entropies were calculated using normal mode analysis (NMA) and plotted against experimental gas-phase entropies for 1665 molecules (blue dots). The parameters could not be generated for the remaining 277 molecules (see Methods). The orange dots represent the positional and orientational entropy as a fraction of the calculated total entropy. (D) The difference in calculated (Y-axis) and experimental (X-axis) entropy of all possible pairs of molecules are plotted as histograms. The area of the circles is proportional to the number of molecule pairs the circle represents. Plots (E) and (F) represent the entropy differences calculated using G4 and NMA, respectively.

Here, we conducted random data sampling with replacement and refitted the GA models for each replicate. We calculated the confidence intervals for the parameters based on 100 bootstrap replicates of the data.

To investigate if different possible conformations of a molecule change the 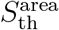 values, i.e., whether we need to include something similar to *S*_conf_ in *S*_vib_, we generated multiple conformations (∼10) of randomly selected 30 molecules (see Methods) and calculated 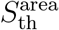 for each of the conformations. The variation of 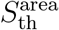 between different conformations was less than 1%. Consequently, in our data set of small molecules, we concluded that 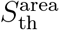 is agnostic to the possible molecular conformations.

### The impact of positive and negative curvature on the entropy

The shape index (𝒮), our definition of surface curvature, value for a perfectly spherical surface is 1, and an inverted concave spherical surface is -1. As surface features move from convex to concave, the 𝒮 value goes from 1 to -1. 𝒮 = 0 represents a saddle, where these two different, positive and negative, curvature meets. In our surface representations of molecules, 0 *<* 𝒮 *<* 1 (𝒮^+^) appears near the surface where two atoms form a bond. Whereas −1 *<* 𝒮 *<* 0 (𝒮^−^) appears where more than two atoms, i.e., more than one patch with 𝒮^+^ overlap. For example, in benzene, 𝒮^+^ are at the C-H bonds, and 𝒮^−^ are at the intersection of two C-H bonds and the center of the carbon-ring (Fig. 2). As expected, coefficients for the deformation terms are negative, as they represent constraints and reduce our MI of the system. Moreover, the absolute coefficient value of the term corresponding to 𝒮^−^ in eqn. 8 (21.46 - Table 1), is substantially higher than that of 𝒮^+^ (6.91 - Table 1), as 𝒮^+^ represents the presence of bonds between two atoms, whereas 𝒮^−^ represents the presence of bonds between multiple and more than two atoms, thus reducing our MI of the system to a great extent.

**Figure 2:**
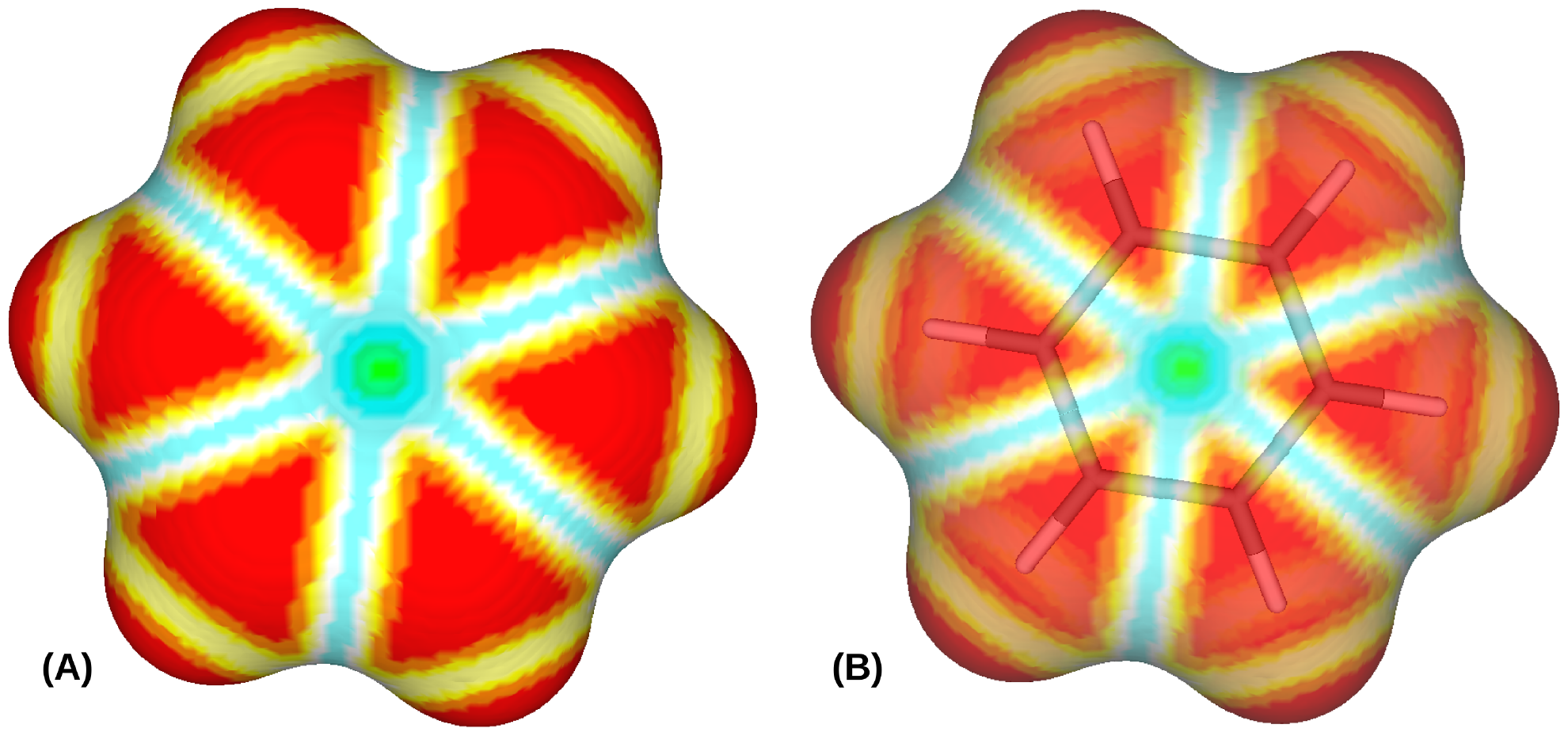
Figure A shows the shape index (see Equation 11) mapped to the Gaussian molecular surface of benzene for *σ* = 0.1. Shape index scales (ranging from -1 to 1) are divided into 9 categories as defined by Koenderink and van Doorn [1992]: (i) 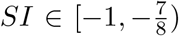 shown in green, (ii) 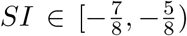 in cyan, (iii) 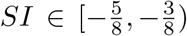 in blue, (iv) 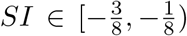 in pale blue (v) 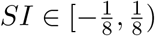 in white, (vi) 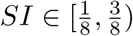 in pale yellow, (vii) 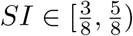 in yellow, (viii) 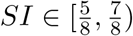 in orange, and (ix) 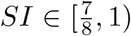 in red. Figure B shows a transparent version of Figure A, along with the embedded model of benzene represented by sticks.

In our datasets, the major fraction of surfaces have 𝒮 value between 0 and 0.99 (61.7%). Another 23.9% of the surface has 𝒮values *>* 0.99, and the remaining 14.4% of the surface has 𝒮 values *<* 0 (Fig. S2 in SI).

### The coefficient of ultraviolet cutoff - connecting *S*_BH_, *S*_entanglement_, and 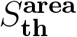

The ultraviolet cutoff, the minimal value of the length used in the derivation of the area law, in eqn. 6 is the *L*_P_, the Planck length. The exact definition of the ultraviolet cutoff, *L*_UV_, will depend on the system under study; for example, if the system under study is a crystal, then *L*_UV_ would be the atomic spacing. The coefficient for the ultraviolet cutoff, *C*_UV_, in eqn. 6 is 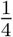 . For a system with *N* coupled harmonic oscillators, Srednicki showed that the *C*_UV_ is 0.30 [Srednicki, 1993], very close to 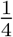 . To calculate the *C*_UV_ in our system, we should change the entropy values from *J/mol* · *K* for 1 mole of molecules to *J/K* for one molecule. The ultraviolet cutoff for our derivation is 1.1 Å, the VDW radius of the hydrogen atom we used. If we ignore the constant term in eqn 8 and focus on the coefficient of the term proportional to the area, we can write,

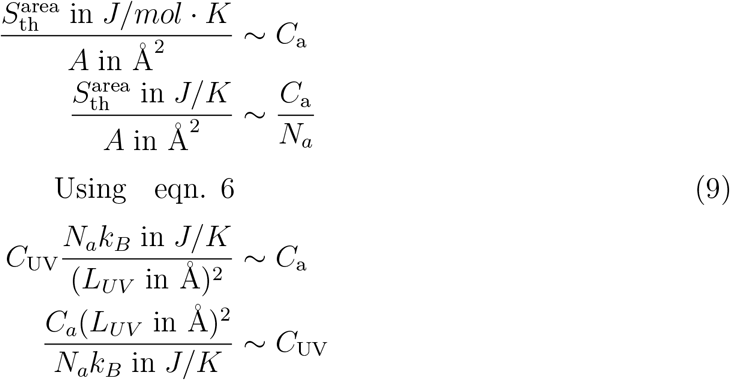

where *A* is the area and *N*_a_ is the Avogadro’s number. The constant *C*_a_ is 3.078 in eqn. 8, and 1.948 *J/K*·Å^2^ if we ignore the deformation terms (Table 1). Putting *L*_UV_ = 1.1 Å, and the *C*_a_ values in eqn. 9, we get *C*_UV_ 0.41 and 0.26, respectively - values very close to the *C*_UV_ values in black hole entropy, and quantum entanglement entropy for *N* coupled harmonic oscillators. Interestingly, for the model where deformation terms are ignored (*S*_area_), the *C*_UV_ is 0.26, or 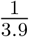, so tantalizingly close to the 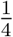 term in black hole entropy, where the wrinkles in the black hole surface are also ignored! Note that the power of *S*_area_ model to predict gas-phase entropy is at par or better than the other currently popular methods compared in this article (Table 1 and Fig. S5 in SI). The area law of entropy, calculated and measured in three different types of complex systems of three vastly different physical dimensions, returns a constant with similar values on all three occasions, thus validating the pertinent idea behind the theory.

### Comparison with entropies calculated using RRHO approximation

RRHO model, which approximates a molecule as a collection of harmonic oscillators, is the most commonly used method to calculate the entropy of a molecule. We curated and calculated entropies, *S*_RRHO-G4_, for 1529 molecules using the quantum-chemical Gaussian-4 (G4) theory (see Methods). We also calculated entropies, *S*_RRHO-NMA_, for 1665 molecules using NMA and molecular mechanics forcefield (see Methods). Both methods show collinearity with the experimental gas-phase entropies for smaller molecules with values less than ≈ 500 *J/mol* · *K* (Fig. 1 B,C). For larger molecules, the methods start deviating from the collinearity (Fig. S4 in SI). RRHO methods use analytical formulas to calculate the positional and orientational entropy (*S*_RR_). For gas-phase entropy less than ≈ 500 *J/mol* · *K, S*_RR_ consists of a significant fraction of the entropy values. As the *S*_RR_ falls below 60% of the total entropy, the deviation from the collinear behavior becomes apparent (Fig. 1 B,C). A possible reason can be that contributions from *S*_anharm_ and *S*_conf_ in *S*_vib_, which the harmonic oscillator approximation cannot capture, increase as the size of a molecule increases. Note that a recent study identifies that conformational entropy accounts for ≈*<* 5% of gas-phase entropy in small molecules [Chan et al., 2021a]. Another possible reason is that the *S*_orie_ is not decoupled from *S*_vib_ for larger molecules, as approximated in the RRHO model.

*S*_RRHO-G4_ and *S*_RRHO-NMA_ values have RMSE values of 24.77 and 45.44 *J/mol* · *K*, respectively, with *S*_Expt_ (Table 1). The values for MAPE for *S*_RRHO-G4_ and *S*_RRHO-NMA_ are 4.09% and 5.41%, respectively (Table 1). The huge difference in RMSE values between *S*_RRHO-G4_ and *S*_RRHO-NMA_ is primarily because *S*_RRHO-G4_ contains the values for smaller molecules, where *S*_RR_ dominates the entropy values. The calculation of *S*_RRHO-G4_ did not converge for larger molecules. Furthermore, in our dataset, the coefficient of determination, *R*^2^, is higher between *S*_RRHO-NMA_ and *S*_Expt_, 0.97, than between *S*_RRHO-G4_ and *S*_Expt_, 0.95 (Table 1). If we include molecules present in all three different sets, 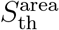, *S*_RRHO-G4_ and *S*_RRHO-NMA_, and calculate RMSE based on those 1326 molecules, the RMSE values are 20.91, 23.38 and 26.21 *J/mol* · *K*, respectively (Table 2. Similarly, for the 1529 molecules common to 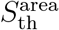 and *S*_RRHO-G4_ sets, RMSE values are 21.47 and 24.76 *J/mol* · *K*, respectively (Table 2). And for the 1665 molecules common to 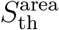 and *S*_RRHO-NMA_ sets, RMSE values are 20.93 and 45.44 *J/mol* · *K*, respectively (Table 2). As the size of the molecules increases, the power of 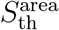 to represent both *S*_RRHO_ and *S*_anharm_ becomes apparent.

**Table 2:** RMSE reported for different combinations based on the data common to the 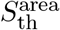, *S*_RRHO-G4_ and *S*_RRHO-NMA_ datasets. Numbers in brackets indicate the number of compounds in common for the groups considered.

| | $S_{th}^{area}$ | $S_{RRHO-G4}$ | $S_{RRHO-NMA}$ |
| --- | --- | --- | --- |
| $S_{th}^{area} \cap S_{RRHO-G4} \cap S_{RRHO-NMA}$ (1326) | 20.91 | 23.38 | 26.21 |
| $S_{th}^{area} \cap S_{RRHO-NMA}$ (1665) | 20.93 | — | 45.44 |
| $S_{th}^{area} \cap S_{RRHO-G4}$ (1529) | 21.47 | 24.76 | — |

### Prediction of relative entropies

Often, relative entropy, i.e., the difference in entropy values between two molecules, is a more helpful quantity than the individual absolute entropy values. To compare the performances of three entropy models in predicting differences in experimental gas-phase entropy values, we calculated the experimental gas-phase entropy difference, δ*S*_Expt_, between all possible pairs of the molecules in a dataset. We compared them with the corresponding difference in the modeled entropy values –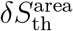, δ*S*_RRHO-G4_, and δ*S*_RRHO-NMA_. The RMSE between the relative entropy values are 30.18, 31.15, and 58.21 *J/mol* · *K* for 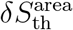, δ*S*_RRHO-G4_, and δ*S*_RRHO-NMA_, respectively.

## 3 Conclusion

Entropy encapsulates missing information (MI) or our ignorance about a system. After decades of theoretical works, since the Bekenstein-Hawkins work on black hole entropy, physicists are converging on that the area of the horizon describes our ignorance or the MI of the matter that’s fallen in — all different ways of internally arranging the building blocks of the black hole to match its outward appearance without knowing what the microstates are. Similarly, for a gas made of molecules, we know the temperature — the average speed of particles — but not what every particle is doing, and the gas’s entropy reflects the MI about the number of options the particles can organize themselves. For two different physical systems of different dimensions, black holes, and entanglement of quantum particles, the area of the systems is proportional to the MI or entropy. We show that proportionality, or the area law, also holds for the thermodynamical entropy of gaseous molecules. The coefficient in the area law in our gas-phase entropy is very close to those in the black hole and quantum entanglement area laws of entropy, indicating the robustness of the underlying idea in three systems of vastly different dimensions.

Calculating thermodynamical entropy using area law allows for calculating molecular entropy faster and more accurately than the currently popular way of approximating the molecules as a collection of harmonic oscillators. In our model, we approximated the atoms to have spherical surfaces with VDW radii. Furthermore, each type of element has been assigned a single VDW value. In molecular mechanics forcefields, used in the calculations of *S*_RRHO-NMA_, the VDW radii depend on the element type and the atom’s local environment. Consequently, the same element can have different VDW radii in different atomic environments. The definition of atomic surface and the VDW radii, possibly considering the atomic environment, can be updated to improve the accuracy of the area law in our model. The speed and accuracy of our method will open up new possibilities for the explicit inclusion of entropy in computational biology methods, such as molecular docking or QSAR (quantitative structure-activity relationships) methods and other methods related to virtual screening.

## 4 Methods

### Data Curation

As the first step, we built a database with experimental gas-phase entropy values (at 25 ° C in *J/mol* · *K*) for various organic compounds (involving elements C, H, N, O, S, P, Cl, Br and I) curated from literature [Guthrie, 2001], [Ghahremanpour et al., 2016], [van der Spoel et al., 2018], [Raychaudhury et al., 2019], [Bains et al., 2022]. Overall, 1942 compounds with corresponding experimental entropies in the range 190 – 1040 *J/mol* · *K* were obtained (see Figure S1 in the Supporting Information). Most of the compounds (≈84%) had entropies below 500 *J/mol* · *K* (Fig. S1 in SI) and had less than five rotatable bonds. High entropies (*>* 700 *J/mol*·*K*) were associated with compounds containing more than 45 heavy atoms and more than ten rotatable bonds.

### Molecular structure calculation

Our curated database contains SMILES strings of chemical compounds. The SMILES strings for each molecule were converted into a single set of 3D coordinates using RDKit [Landrum, 2020]. The atoms’ van der Waals (VDW) radii were assigned using the software OpenBabel [O’Boyle et al., 2011]. Note that the VDW assignment by OpenBabel does not depend on the local environment. For example, all carbon atoms will have the same VDW radius; no further parameterization is needed for individual small molecules. The structures were minimized using the Universal Force Field [Rappe et al., 1992] (UFF) implemented in RDKit.

To understand the impact of conformational variability on 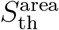, we generated multiple conformers using RDKIT [Ebejer et al., 2012]. For some 30 randomly selected molecules (with more than three rotatable bonds), up to ten conformers were generated using RDKIT. Since the generated conformers can be structurally similar, only conformations at least 0.50Å root mean square deviation (RMSD) apart from one another were retained. As a result of this filtering, the final tally of generated conformers was for some compounds less than 10. Each conformation was then subjected to geometry optimization using the AM1[Dewar et al., 1985] Hamiltonian in the semi-empirical program MOPAC[Stewart, 2016].

### Theoretical Entropy Calculation

To calculate the gas phase entropies for the molecules, the quantum-chemical Gaussian-4[Curtiss et al., 2007] (G4) level of theory was used. The G4 theory has been shown to provide a good compromise for thermochemistry calculations in comparison to the other methods tested (see Ghahremanpour et al. [2016] and Ghahremanpour et al. [2018]). The G4 entropies for almost 1000 compounds were taken from the Alexandria library [Ghahremanpour et al., 2018]. Entropies for an additional ∼500 compounds were computed in-house. Compounds with convergence issues and those that took more than 24 hours of computing time were excluded. The calculations were performed using Gaussian 09[Frisch et al., 2009]. The OpenBabel tool *obthermo* was subsequently used to extract thermochemistry data from Gaussian output files.

We also calculated entropy values estimated using normal-mode analysis (NMA), *S*_RRHO-NMA_ for 1665 molecules. Here, The structures of all compounds were first energy-minimized using GROMACS [Abraham et al., 2023]. Normal mode analysis was then carried out using GROMACS, and the entropy estimates were obtained using the “gmx nmeig” module.

### Molecular surface calculation

In this study, the surface is defined analytically as *M* = *G*^−1^(**x**)|_c_ where the function *G* is a weighted sum of atom-centered Gaussians [Gabdoulline and Wade, 1996] given by:

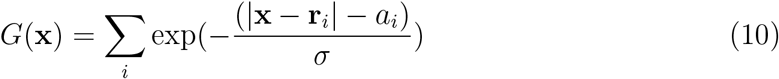

where *r*_i_ is the position of the *i*^th^ atom centre and *a*_i_ the corresponding VDW radius. The surface *M* is defined by values of **x** for which the function *G*^−1^(**x**) = *c*. The smoothing parameter 0 ≤ *σ* ≤ 1 affects the level of detail associated with the surface. Larger values of *σ* smooth out the surface details, while the smaller values of *σ* preserve more details of the surface feature. Please see Figure S3 in the SI to compare the impact of different *σ* on the level of surface detail. The Gaussian surface isovalue *c* controls the volume enclosed by the surface. In this article, we calculated the surfaces at the isovalue *c* = 1.0 and used a smoothing factor (*σ*) of 0.1. Using in-house software, the molecular surfaces were generated using the Marching cubes algorithm [Vega et al., 2019].

### Surface curvature

The VDW surface for the molecules defined by Equation 10 is a continuously differentiable function and is defined analytically at every point. From the analytical expression of the surface, we can calculate principal curvatures at every point. If we can draw a small normal plane at a point on a surface and calculate the curvature of every line going through the point, then the highest and lowest curvatures of those lines, calculated at the point, are the principal curvatures *κ*_1_ and *κ*_2_ (*κ*_1_ *> κ*_2_) of the surface at that point. The principal curvatures measure how the surface bends in different directions at a point (see do Carmo [1976]) and can be calculated analytically from the first- and second-order partial derivatives of the surface function [Goldman, 2005, Xia et al., 2014]. The principal curvatures can be further combined to define the shape index (𝒮) [Koenderink and van Doorn, 1992]:

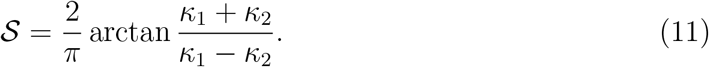

The shape index (𝒮) measures the local shape of a surface and varies from -1 (concave) to 1 (convex).

### Shape index probabilities

The molecular surface, once calculated, is discretized by a triangle mesh using the Marching cubes algorithm, i.e., covering the surface with triangles. After triangulation, the shape index values are calculated at the center of each mesh triangle. The probabilities *p*^±^ that a specific surface patch of a molecule will have a shape index value of 𝒮^±^ are determined by binning the shape index values. We should emphasize that we calculate each molecule’s *p*^±^ values independently. The entropy calculation using equation 8 can be impacted by the bin width and the smoothing factor of the Gaussian surface function. We varied the bin numbers between 64, 128, and 256 and the smoothing factor *σ* between 0.1, 0.3, and 0.5. We then calculated the Pearson correlation coefficient between the shape entropies and the experimental gas-phase entropy to determine the effect of *σ* and the number of bins (see Table S1 in the SI). While the smoothing factor *σ* affects the correlations to some extent, the bin width or number has little to no impact. We chose 64 bins to calculate the histograms and the probability density of the curvedness values using *σ* = 0.1 for the rest of the work.

### Data Fitting

To identify suitable values of the parameters (*S*0, *a, b, c*) in Equation 8, we used a genetic algorithm (GA). In the GA search, the upper and lower bounds for the parameters were set to ±200, ±100, ±50, ±50, while the population size was varied between 100 and 250. Crossover and mutation probabilities were set to 0.75 and 0.25, respectively. The fitness function to be maximized was set to the inverse of the root mean square error and ran the algorithm for 100 cycles. The calculations were carried out using the GA [Scrucca, 2013, 2017] package in R.

## Data Availability

All data used in this study is available in the Supporting Information.

## Software Availability

The gas phase entropy prediction is available online at https://vvishwesh.github,io/surfent.

## Supporting Information Available

- An Excel file with the observed and predicted gas-phase entropies for the different compounds (in SMILES format), including gas-phase entropy calculated using NMA with the molecular mechanics force fields GAFF-ESP and GAFF-BCC.
- A brief introduction to von Neumann entropy and quantum entanglement entropy.
- Figure S1 showing the statistics of curated data and Figure S2 showing the distribution of shape index values 𝒮 in our dataset.
- Figure S3 showing the effects of smoothing parameter *σ* and Table S1 showing the effect number of bins for calculating histogram on the shape entropy.
- Figure S4 showing the performance of *S*_RRHO-G4_ and *S*_RRHO-NMA_ as a function of number of atoms, and Figure S5 showing the performance of *S*_area_ in predicting *S*_Expt_.

## Acknowledgements

VV acknowledges financial support from the Research Council of Norway (Grant No. 262152). The authors thank Dr. Sudip Roy for his valuable input to the manuscript.

## Author Contributions

VV and AR conceived the study. All authors wrote the manuscript collaboratively.

## Competing interests

The authors declare no competing interests.

## Notes

### Competing Interest Statement

The authors have declared no competing interest.

