## Supporting Information for "The Area-law of Molecular Entropy: Moving beyond Harmonic Approximation"

#### Quantum mechanical entropy

In quantum mechanics, the state of a system S is usually represented by a state vector $|\psi\rangle$, which is an element of a Hilbert space $\mathcal{H}_{\rm S}$. But it is possible that there is some uncertainty associated with exactly which state the system is in and that the probability that the system is in state $|\psi_i\rangle$ is given by $p_i$. Under such circumstances the quantum state of the system is *not* given by a "pure" state $|\psi\rangle$ but by the density operator $\rho = \sum_i p_i |\psi_i\rangle \langle\psi_i|$. The uncertainty related to which state the system is in is measured by the quantum entropy or von Neumann entropy

$$S_{\rm vN} = -{\rm Tr}\left[\rho \ln \rho\right]. \tag{S1}$$

For the pure state $|\psi\rangle$ the density operator is $\rho = |\psi\rangle \langle\psi|$ and the von Neumann entropy is defined to be zero. Note that $\rho$ in some given basis need not be diagonal but when it is its von Neumann entropy is equivalent to the classical form of Shannon entropy.

Whether the von Neumann entropy corresponds to thermodynamic entropy is still an active research area and outside the scope of this article. If, however, we construct the density matrix following Jaynes's argument jaynes1957information2, an argument similar to deriving thermodynamic entropy from information theory, as mentioned earlier, the connection between the quantum and thermodynamic entropies becomes apparent.

Briefly, to build a density matrix $\rho$ we need to know the probability $p_i$ for the system to be in the state $|\psi_i\rangle$, information that is not readily available. However, according to Jaynes, if we have access to measured quantities, for example, the average energy $\langle E|E\rangle = {\rm Tr}[\rho\hat{H}]$, where $\hat{H}$ is the Hamiltonian operator, then one can construct the density matrix by maximizing the entropy function subject to the constraints that the expectation values computed out of the density matrix yield the observed measurements. In the particular case where we impose the constraint that the average energy $\langle E|E\rangle$ is constant, one obtains the Gibbs state as the density matrix $\rho$.

#### Quantum entanglement entropy

In quantum mechanics, microstates are described, in general, by a density matrix. However, a non-zero quantum von Neumann entropy is associated with this state even if it originates in a pure state when part of the system is inaccessible to the observer. In the quantum world, the state of two or more particles can connect so that observations on each can correlate. They cannot be described independently of the others. This phenomenon is called quantum entanglement. Let us consider a bi-partite quantum system consisting of A and B subsystems. The joint state $|\psi\rangle_{\rm AB}$ of the combined system must be an element of the Hilbert space $\mathcal{H}_{\rm AB}$. This Hilbert space has a tensor product structure $\mathcal{H}_{\rm AB} = \mathcal{H}_{\rm A} \otimes \mathcal{H}_{\rm B}$, where $\mathcal{H}_{\rm A(B)}$ is the Hilbert space of the subsystem A(B). If the state $|\psi\rangle_{\rm AB}$ of the combined system is entangled it means that it *cannot* be written as $|\psi\rangle_{\rm A} \otimes |\psi\rangle_{\rm B}$. For example, two spin-half particles can form the single state $\frac{1}{\sqrt{2}}\left(|\uparrow\rangle \otimes |\downarrow\rangle - |\downarrow\rangle \otimes |\uparrow\rangle\right)$ which is an entangled state.

Now suppose the state of the combined system is given by an entangled state $|\psi\rangle_{\rm AB}$ and that an observer, say Alice, wants to measure a property of the quantum system A, but she doesn't want to disturb or doesn't have access to system B. Then the state of A that she must work with is *necessarily* given by a mixed state described by the *reduced* density matrix $\rho_{\rm A} = {\rm Tr}_{\rm B}\, \rho_{\rm AB}$, where $\rho_{\rm AB} = |\psi\rangle_{\rm AB} \langle\psi|_{\rm AB}$ and the trace is taken *only* over the Hilbert space of system B reflecting Alice's ignorance about the second subsystem. The combined system AB is at zero temperature, which is implied by the fact that the state of the combined system is the pure state $|\psi\rangle_{\rm AB}$. Otherwise, we would have to describe the state of the combined system in terms of a density matrix known as the Gibbs state. Thus, despite the zero temperature of the combined system, the von Neumann entropy $S_{\rm vN}$ of $\rho_{\rm A}$ is non-zero due to non-trivial entanglement. The positive entropy that arises from the measurement of a subsystem is called entanglement entropy, which is unavoidable in any quantum measurement. The entanglement entropy depends on the area of the system under observation, as discussed in the main text.

### Dataset

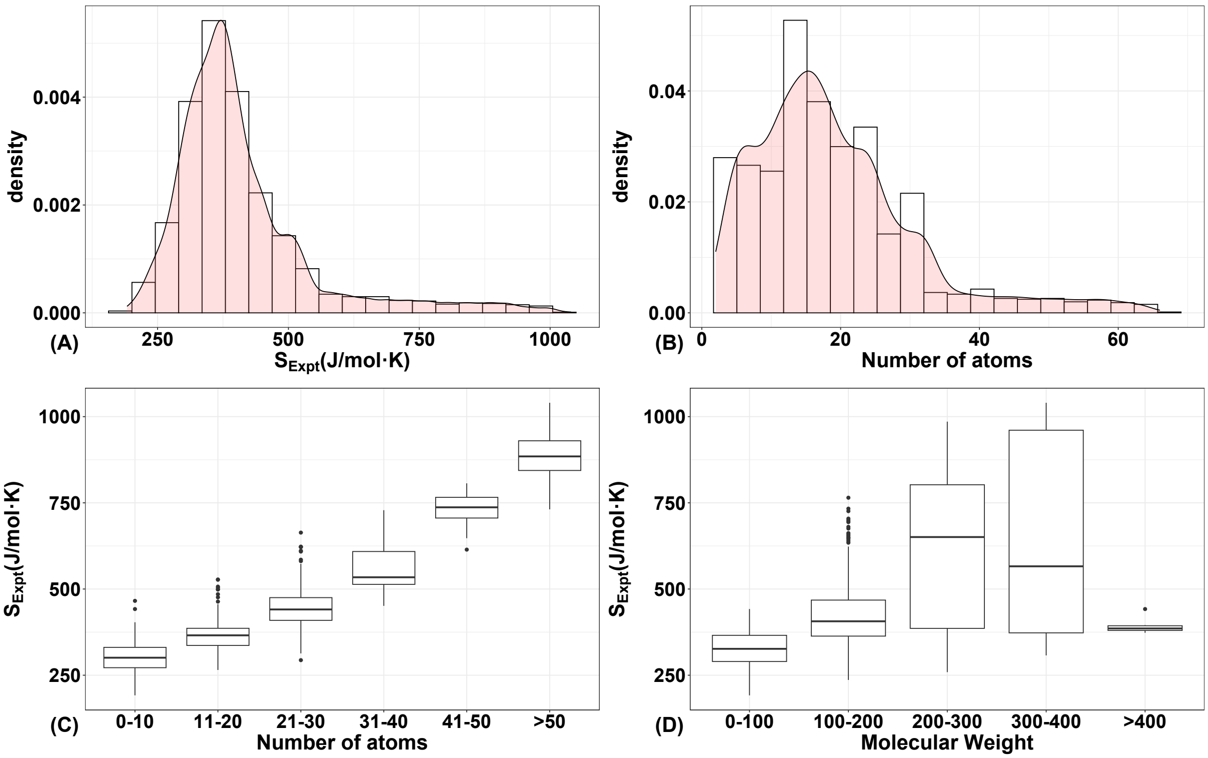


**Figure S1:** The figure provides a visual summary of the gas phase entropies for a set 1942 organic molecules. (A) Histogram distribution of the entropies (B) (A) Histogram distribution of the number of atoms (C) Boxplot showing that entropies typically increase with the number of atoms in the molecule. (D) Boxplot showing the variation of the entropies with the molecular weight

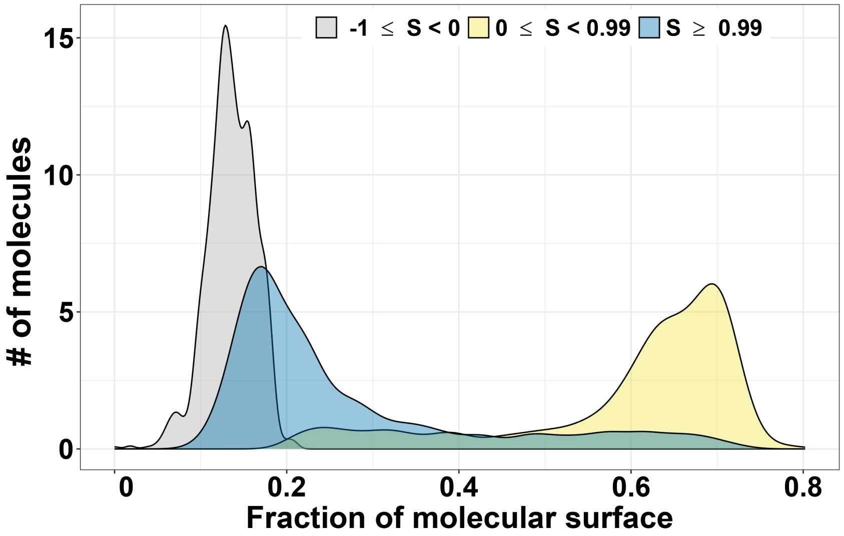


**Figure S2:** Distribution of $\mathcal{S}$ values in our dataset of 1942 molecules. The major fraction, 61.7%, of surfaces have $\mathcal{S}$ value between 0 and 0.99 (yellow). Another 23.9% of the surface has $\mathcal{S}$ values $> 0.99$ (blue), and the remaining 14.4% of the surface has $\mathcal{S}$ values $< 0$ (gray).

#### Parameters

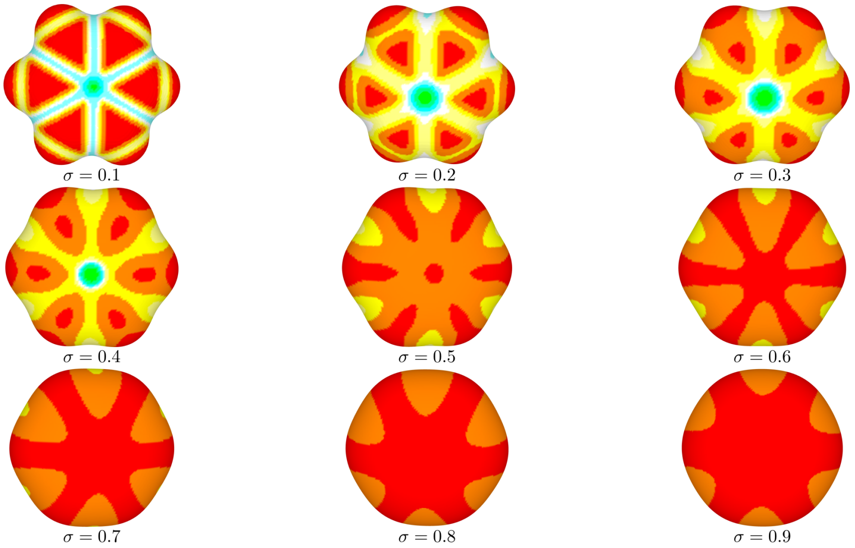


**Figure S3:** Impact of the surface smoothing factor $\sigma$ on the level of detail in the surface. Larger values of $\sigma$ smooth out the details of the surface, while the smaller values of $\sigma$ preserve more details of the surface features.

| $\sigma$ | **#Bins** | **RMSE in** $J/mol \cdot K$ |
|---|---|---|
| 0.1 | 64 | 21.33 |
| 0.1 | 128 | 21.79 |
| 0.1 | 256 | 22.79 |

**Table S1:** Table shows the impact of the surface smoothing factor $\sigma$ and the number of bins on the root mean square error.

### Results

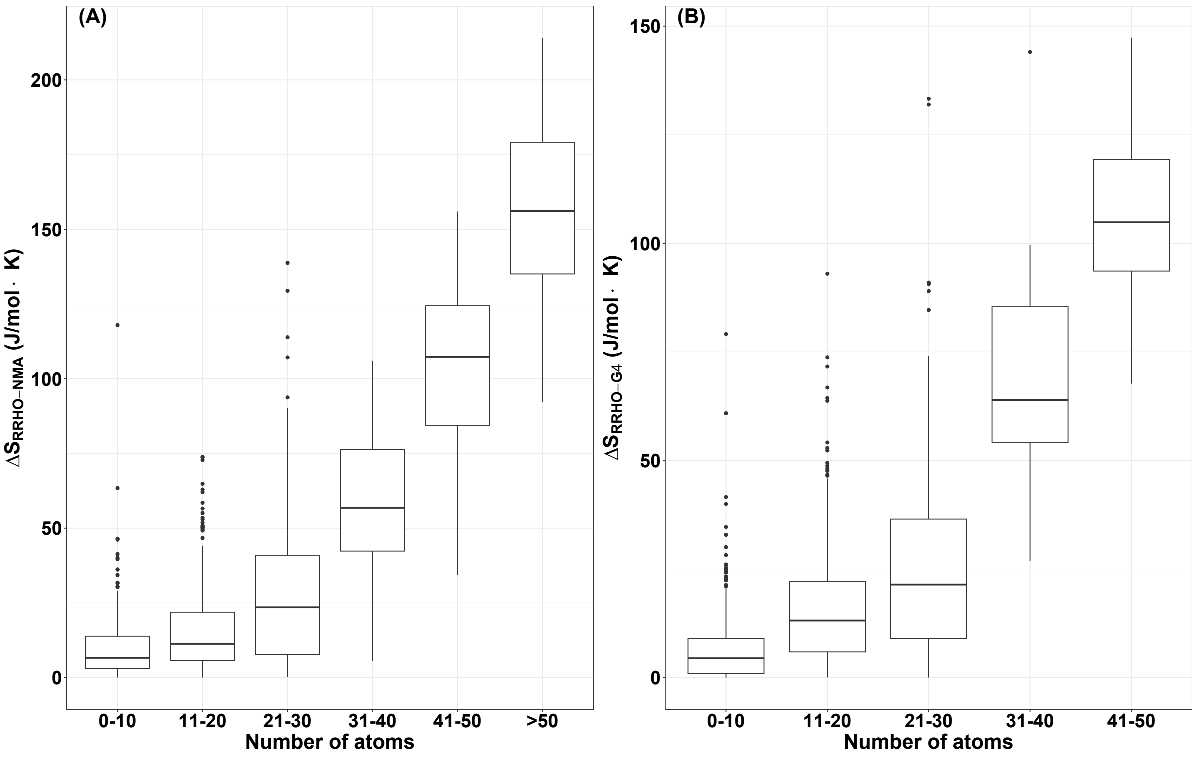


**Figure S4:** Box plot of the absolute deviations between the $S_{\text{Expt}}$ and (A) $S_{\text{RRHO-NMA}}$ (B) $S_{\text{RRHO-G4}}$ entropies. In both cases, larger deviations (>50 $J/mol \cdot K$) are typically associated with molecules containing more than 30 atoms.

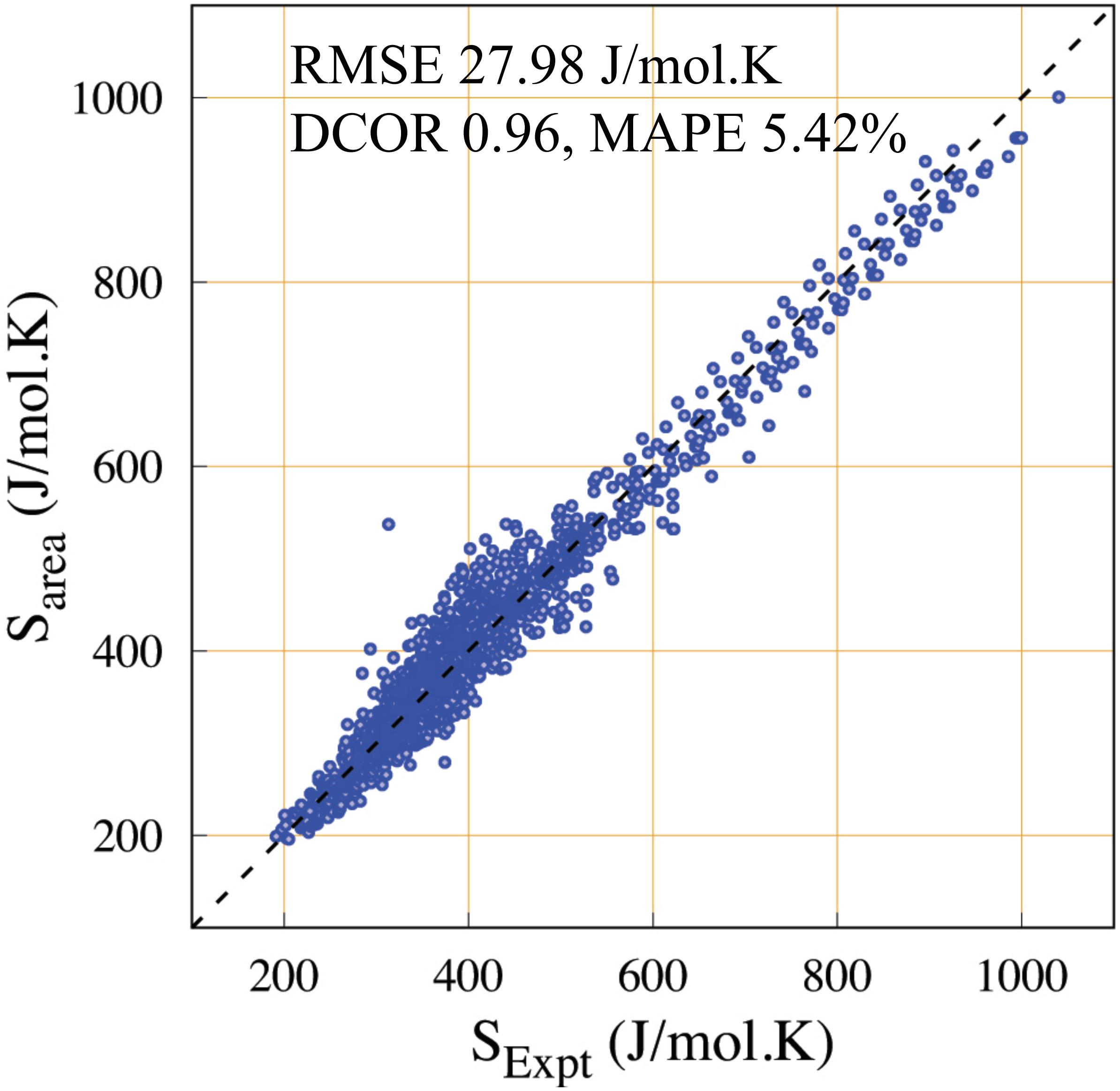


**Figure S5:** $S_{\text{area}}$, the thermodynamical entropies calculated from the area law (Table 1 in the main text), without considering the effect of surface deformations, are plotted against experimental gas-phase entropies for 1942 molecules. The root mean square error (RMSE) between the calculated and experimental entropy is 27.98 $J/mol \cdot K$. The correlation (DCOR) between the values, calculated using distance correlation (see Methods), is 0.96, and the mean average percentage error (MAPE) is 5.42%. The dotted line represents the line where the values of the X and Y axes are equal.

#### References
